# NLRP3 inflammasome is regulated in osteoclasts through a Tmem178-dependent mechanism that restricts calcium influx

**DOI:** 10.1101/2025.07.28.667255

**Authors:** Khushpreet Kaur, Yael Alippe, Chun Wang, Nicholas P. Semenkovich, Saumya Bhagat, Kunjan Khanna, Yongjia Li, Nitin Pokhrel, Timothy Peterson, Deborah J. Veis, Yousef Abu-Amer, Roberta Faccio, Gabriel Mbalaviele

## Abstract

Osteoclasts (OCs) differentiate from macrophages in response to RANKL. Here, we investigated the role of the NLRP3 inflammasome in mouse macrophages, with or without exposure to RANKL. Unexpectedly, we found that NLRP3 expression gradually declined during osteoclastogenesis but could be restored with LPS treatment. LPS and nigericin robustly activated this inflammasome in macrophages, as expected, but not in OCs. Mechanistically, we identified Tmem178, a protein that restrains Ca^2+^ release from the endoplasmic reticulum (ER) and highly expressed in OCs, as an inhibitor of this inflammasome. Notably, NLRP3 inflammasome activation was robust in OCs lacking Tmem178 or wild-type (WT) OCs exposed to high calcium concentrations. *In vivo* studies demonstrated that under the conditions where OCs efficiently release Ca^2+^ from bone, inflammasome formation was enhanced. Furthermore, deletion of *Nlrp3* rescued osteopenia in *Tmem178^−/−^* mice. Thus, we found that Tmem178 uniquely restricts Ca^2+^ release from ER in OCs, thereby suppressing NLRP3 inflammasome activation.

**One Sentence Summary:** The NLRP3 inflammasome is silenced in the OC lineage by Tmem178 to prevent pathological bone loss.

## INTRODUCTION

Various organisms operate as complex but integrated systems in which specialized proteins act as sentinels, detecting and responding to threats and danger signals to maintain tissue integrity and homeostasis. Among these guard mechanisms, the inflammasome sensor NLRP3 stands out as a critical detector of both pathogen-associated molecular patterns (PAMPs) and sterile damage-associated molecular patterns (DAMPs) ^1–3^. Activation of the NLRP3 inflammasome triggers the release of IL-1β and IL-18 and can lead to pyroptosis^4,5^. While activation of the NLRP3 inflammasome is essential for maintaining tissue homeostasis in response to infection or injury, its dysregulation has been linked to chronic inflammation and various pathological conditions ^6–10^.

The NLRP3 inflammasome plays a key role in bone loss across a wide range of conditions, from low-grade inflammation (e.g., aging, experimental hyperparathyroidism, and menopause) to high-grade inflammatory states (e.g., autoinflammatory disorders, periodontitis, and osteomyelitis) ^6,7,11–14^. Its ability to cause bone resorption in various diseases may stem from both direct (autonomous) and indirect (non-autonomous) effects on OC lineage cells (hereafter referred to as OCs for simplicity). The non-autonomous actions include the release of osteoclastogenic factors by immune cells, osteoblasts, osteocytes, and other stromal cells in the bone microenvironment ^15–17^. These mediators, particularly IL-1β, can act on neighboring cells, directly and indirectly stimulating OC activity and bone resorption ^8,14,18,19^. While non-autonomous actions of the NLRP3 inflammasome in bone are well established, its autonomous role in OCs remains less understood.

OCs exhibit finely tuned calcium oscillations that are essential for their differentiation and resorptive function ^20–22^. Localized calcium puffs occur at acid-secreting sites of bone-attached OCs, which further regulate their resorptive activity ^23,24^. Intracellular calcium is primarily stored in organelles such as the lysosomes, endoplasmic reticulum (ER), and mitochondria, where it is tightly regulated ^25^. Transmembrane protein 178 (Tmem178), a key protein located on the ER membrane, plays a crucial role in controlling cytosolic calcium levels by restricting the release of calcium from the ER into the cytoplasm ^26,27^. Given the critical role of Ca^2+^ in activating the NLRP3 inflammasome ^26,28–31^ and the reported low bone mass phenotype of Tmem178 deficient mice, we hypothesized that the interplay between this calcium regulator and the NLRP3 inflammasome regulates OC biology.

In this study, we found that while the NLRP3 inflammasome is activated in BMDMs, it remains silenced in the OC lineage unless calcium influx is pharmacologically enhanced (e.g., through calcium supplementation or release from bone matrix) or following genetic ablation of Tmem178.

## RESULTS

### The NLRP3 inflammasome pathway is silenced *in vitro* in OCs

The NLRP3 inflammasome plays a critical role in bone resorption, as its product, IL-1β, is a potent stimulator of OC activity ^3,12,15^. However, the intrinsic actions of the NLRP3 inflammasome in OCs remain poorly understood. To address this knowledge gap, we analyzed NLRP3 expression in BMDMs treated with RANKL for 4-5 days to induce OC differentiation. Immunoblotting and qRT-PCR analyses revealed a gradual decrease in NLRP3 expression in OC cultures, which was dependent on the duration of RANKL exposure (Fig. 1A-C). This decline was reversed by treatment with lipopolysaccharide (LPS), a known inducer of NLRP3 expression (Fig. 1B, C). Notably, LPS and nigericin, but not the stimulus alone, induced the secretion of IL-1β, the release of lactate dehydrogenase (LDH) - a marker of pyroptosis, as it is released only upon plasma membrane rupture - and the cleavage of GSDMD. These responses were all reduced in the OC cultures compared to BMDM cultures (Fig. 1D-F; Fig. S1).

**Fig. 1.**
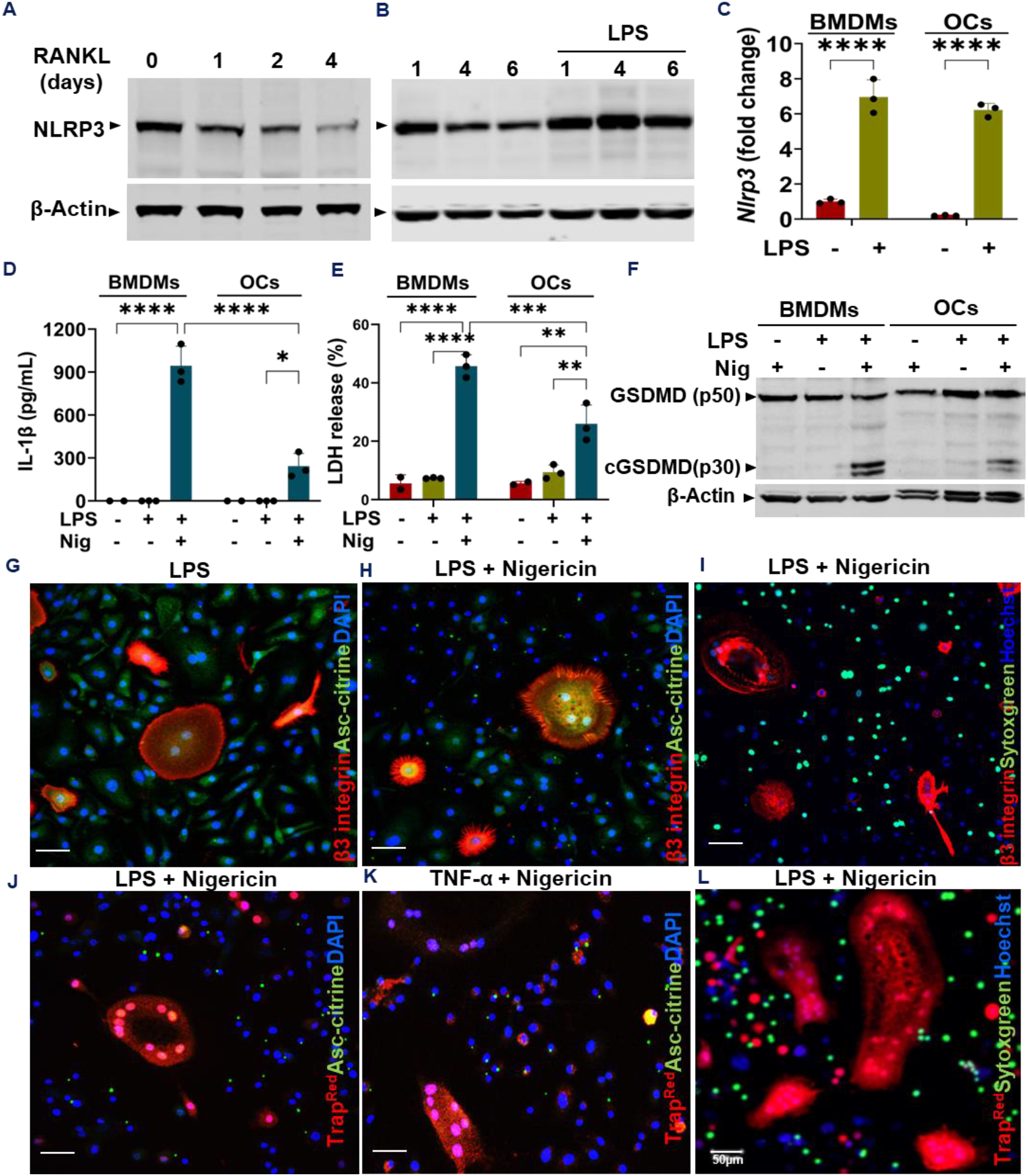
The NLRP3 inflammasome pathway is silenced in OCs. BMDMs from WT (A-F, I), *Asc-citrine* (G, H); and *Trap*^*Red*^;*Asc-citrine* mice (J-L) were treated with RANKL for 5 days or remained RANKL naïve. Cells were primed with 100 ng/ml LPS (B-J;L) or 10 ng/ml TNF-α (K) for 3 h and stimulated with 15 µM nigericin for 45 mins (D-F; H-L). Samples were analyzed for NLRP3, GSDMD, cleaved GSDMD (cGSDMD), and β-Actin by immunoblotting (A,B,F) or for *Nlrp3* expression by qRT-PCR (C). IL-1β (D) and LDH (E) levels in the supernatants were measured by ELISA and the cytotoxicity assay kit, respectively. Assessment of ASC-speck formation (G,H,J,K) or Sytox green uptake by live cells was imaged using a confocal microscope (I, L). Scale bar 50 µm. 2-way ANOVA with Tukey’s multiple comparison test. *p<0.05, **p<0.005; ***p<0.001; ****p<0.0001

To identify cells competent in forming the NLRP3 inflammasome within the heterogeneous OC cultures, we analyzed *Asc-citrine* BMDMs treated with RANKL for the expression of β3 integrin, a marker of OCs. LPS alone did not induce the formation of ASC specks and the uptake of the cell-impermeant DNA-binding dye, Sytox green, in either β3 integrin-negative (i.e., BMDMs) or β3 integrin-positive (i.e., OCs) cells (Fig. 1G, Fig. S2A, B). Unexpectedly, only BMDMs, but not OCs, were ASC speck-positive and showed Sytox green uptake in response to LPS and nigericin treatment (Fig. 1H, I). Similarly, LPS and nigericin failed to induce the formation of ASC specks (Fig. S2C) and Sytox green uptake (Fig. S2D) in OCs expressing constitutively activated NLRP3 ^32^. To investigate whether the inability to nucleate the NLRP3 inflammasome complex occurred early during osteoclastogenesis, we analyzed this response in cells expressing tartrate-resistant acid phosphatase (TRAP), which is expressed earlier than β3 integrin. For this, we used BMDMs from *Trap*^*Red*^; *Asc-citrine* mice, in which td-tomato expression is driven by the *Acp5* promoter ^33,34^. In these cells, LPS and nigericin-induced ASC speck formation was observed only in *Trap*^*Red*^-negative cells, not in *Trap*^*Red*^-positive cells (Fig. 1J). Likewise, treatment with TNF-α and nigericin induced speck formation in BMDMs but not in OCs (Fig. 1K). Consistent with the ASC speck data, LPS and nigericin treatment induced Sytox green uptake in BMDMs but not in β3 integrin-positive OCs (Fig. 1L). Collectively, these data suggest that as soon as BMDMs commit to the OC lineage, they lose the ability to form the NLRP3 inflammasome complex under standard culture conditions.

### OCs assemble inflammasomes *in vivo*

To assess the status of NLRP3 inflammasome activation in OCs *in vivo, Trap^Red^*; *Asc-citrine* mice were subcutaneously injected over the periosteum of the calvaria with PBS or RANKL for 5 consecutive days. Additionally, some RANKL-treated mice were inoculated with LPS for 6 h (Fig. 2A). Confocal microscopy of cryopreserved specimens proximal to the calvaria sutures revealed the presence of ASC specks inside cells embedded in bone, presumably osteocytes, as well as in mononucleated cells and multinucleated TRAP^+^ OCs inside the bone marrow cavity (Fig. 2B; Fig. S3). Although ASC specks were observed in PBS-treated mice, their numbers were significantly higher in mice treated with RANKL in the absence or presence of LPS (Fig. 2C). These results suggest that OCs are capable of assembling inflammasomes *in vivo*, a process that is enhanced under conditions of high bone turnover, such as those induced by RANKL or LPS.

**Fig. 2.**
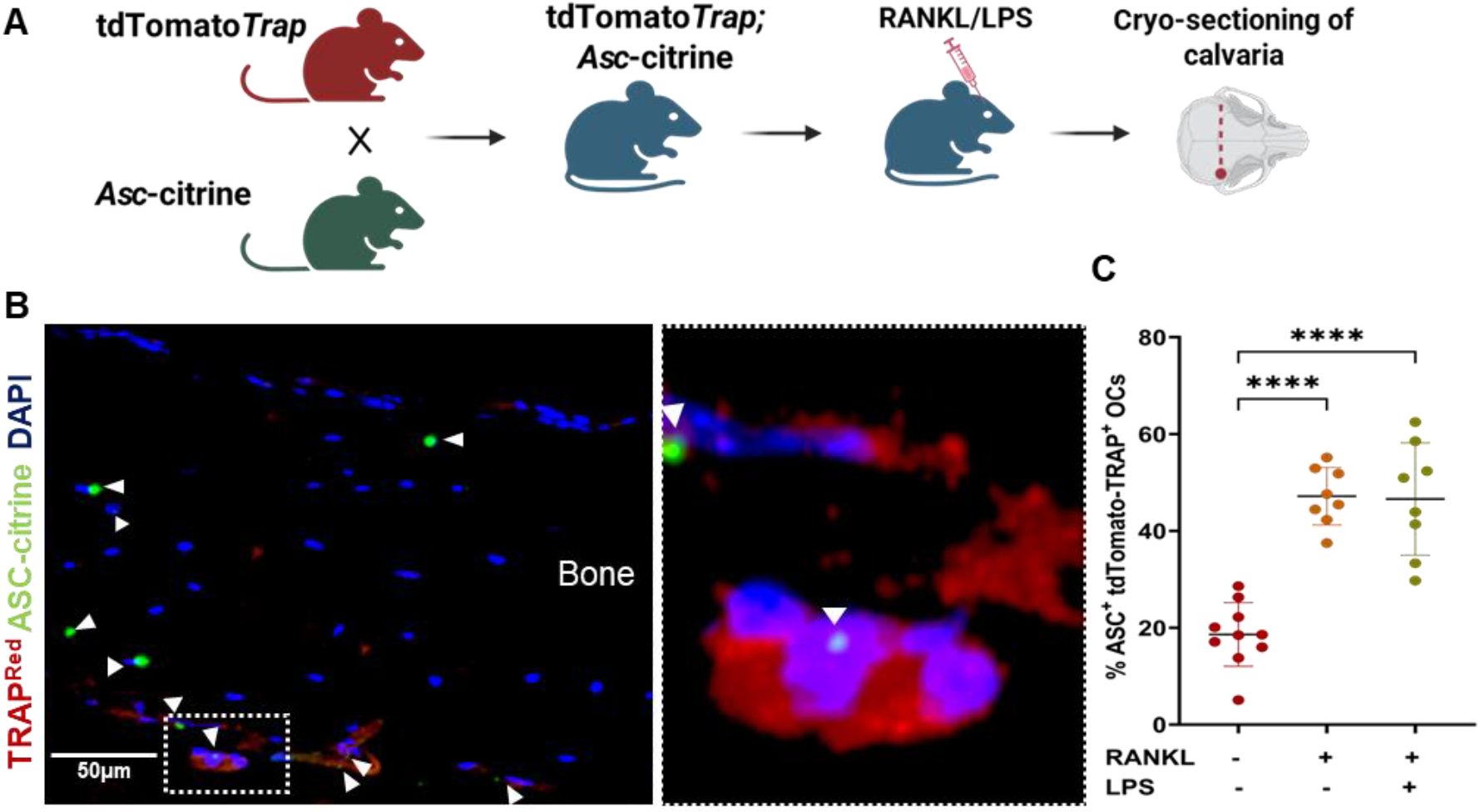
OCs assemble NLRP3 inflammasome *in vivo*. *Trap*^*Red*^;*Asc-citrine* mice were injected with RANKL over the calvarial periosteum for 5 days, then with LPS for 6 h prior to harvest (A). Histological sections showing ASC specks (green dots) in TRAP^Red^ OCs (B), and quantitative data (C). Scale bar: 50 µm. Data were analyzed by 1-way ANOVA, ****p<0.0001.

### Tmem178 inhibits the activation of the NLRP3 inflammasome in OCs

Since the NLRP3 inflammasome could be activated in OCs *in vivo* but not *in vitro*, we aimed to elucidate the mechanisms underlying this differential activation. We performed RNA-seq profiling of BMDMs that were either untreated or treated with RANKL for 5 days, followed by exposure or no exposure to LPS for 3 h. Among the differentially expressed genes (Fig. S4A), *Tmem178* stood out as being markedly upregulated in OCs compared to BMDMs (Fig. S4B). Previous studies have also demonstrated that *Tmem178* was overexpressed in OCs compared to BMDMs, and negatively regulated IL-1β secretion by BMDMs through inhibition of the NLRP3 inflammasome ^26,35^. Consistent with these findings, RANKL treatment induced Tmem178 mRNA expression, which was further enhanced upon cell exposure to LPS (Fig. 3A). Given the cellular heterogeneity of OC cultures, additional steps were taken to separate mononucleated cells (e.g., undifferentiated BMDMs and OC precursors) from multinucleated cells (e.g., mature OCs). The results showed that multinucleated OCs expressed higher levels of Tmem178 compared to mononucleated cells, both in the absence and presence of LPS (Fig. 3B).

**Fig. 3.**
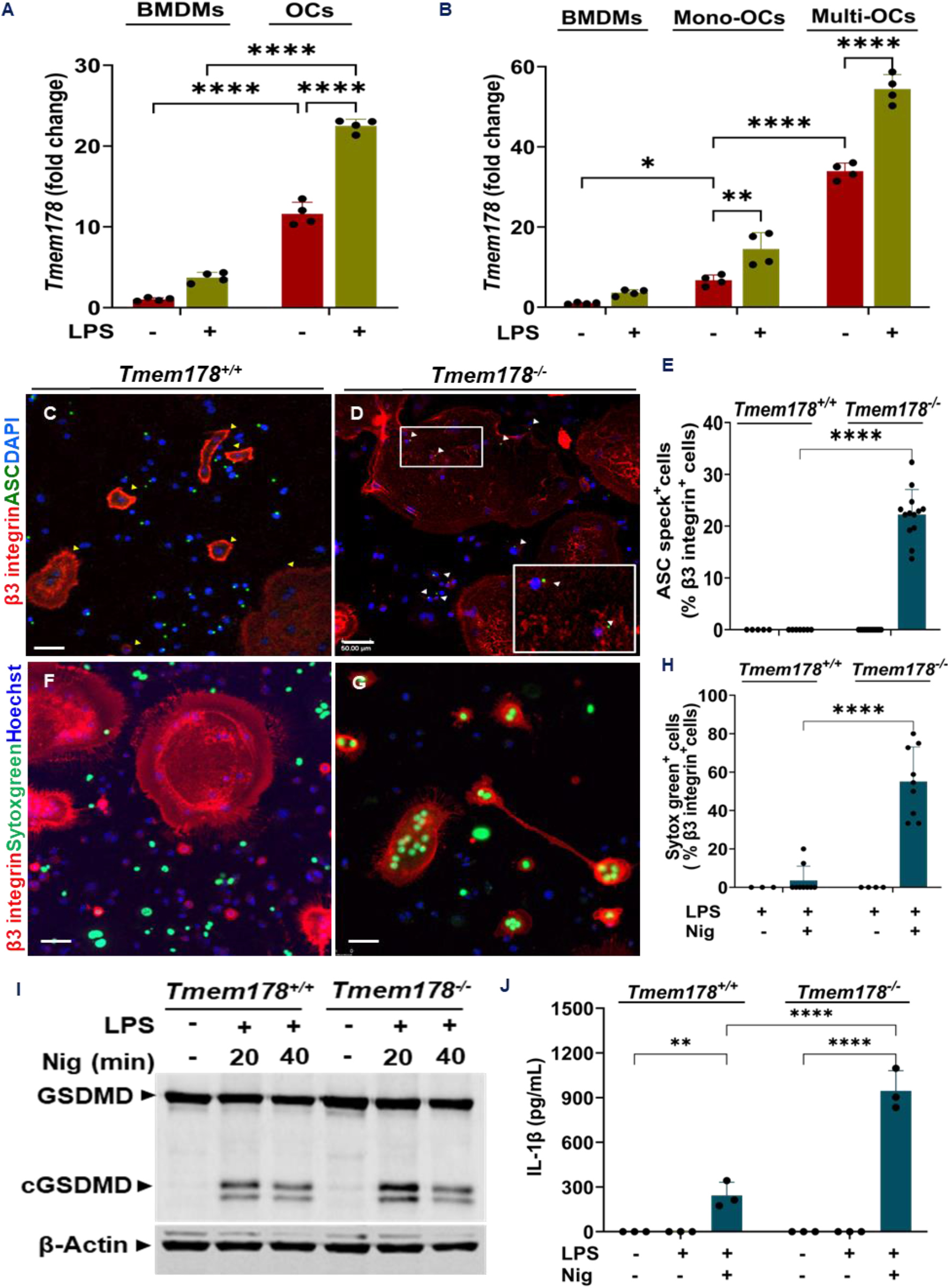
Tmem178 inhibits the activation of the NLRP3 inflammasome in the OC lineage. qRT-PCR analysis of *Tmem178* expression in BMDM and unfractionated OC cultures (A), BMDMs vs OC culture separated into mononuclear and multinuclear fractions (B) were treated with or without LPS for 3 h. *Tmem178*^+/+^ and *Tmem178^−/−^* were treated with LPS for 3 h and nigericin for 45 min (C-J). Cells were stained with β3 integrin antibody (red), followed by fixation and staining using ASC antibody (green), and counterstained with DAPI/Hoechst (blue). The yellow arrow indicates β3 integrin-stained OCs, and white arrowheads indicate ASC specks (C, D). RANKL-treated cells exposed to LPS were incubated with Hoechst followed by nigericin (nig) and Sytox green for 45 min to assess the Sytox green uptake through GSDMD pores (F, G), and quantitative data (E, H). Immunoblotting for GSDMD and cGSDMD in cell lysates after exposure to LPS and nigericin for 20 and 40 min (I). Secreted IL-1β levels measured in enriched multinucleated OC’s culture supernatants by ELISA (J). Scale bar 50 µm. 2-way ANOVA with Tukey’s multiple comparison test, *p<0.05; **p<0.005; ****p<0.0001.

To investigate the role that Tmem178 plays in NLRP3 inflammasome activation in OCs, *Tmem178*^+/+^ and *Tmem178^−/−^* OC cultures were treated with LPS and nigericin, followed by immunostaining with ASC and β3 integrin antibodies, and counterstaining with DAPI. While ASC specks were exclusively observed in *Tmem178*^+/+^ BMDMs (Fig. 3C), they were present in *Tmem178^−/−^* BMDMs and OCs (Fig. 3D, E; Fig. S5). Likewise, Sytox green uptake, which occurred only in *Tmem178*^+/+^ BMDMs but not *Tmem178*^+/+^ OCs (Fig. 3F; Fig. S6A), was observed in *Tmem178^−/−^* BMDMs and OCs (Fig. 3G, H; Fig. S5B). This response was abolished in *Tmem178^−/−^*;*Nlrp3^−/−^* cells (Fig. S7A). Furthermore, GSDMD cleavage (Fig. 3I) and IL-1β secretion (Fig. 3J) induced by LPS and nigericin were higher in *Tem178^−/−^*OCs compared to their *Tmem178*^+/+^counterparts. Collectively, these findings suggest that the calcium gatekeeper, Tmem178, functions as an inhibitor of the NLRP3 inflammasome, not only in macrophages but also in the OCs.

### Tmem178 loss triggers NLRP3 inflammasome activation via increased intracellular calcium levels

Tmem178 resides in the ER, where it binds to stromal interaction molecule 1 (Stim1) and inhibits the activation of store-operated calcium entry (SOCE), thereby preventing increases in cytoplasmic Ca^2+^, an important activator of the NLRP3 inflammasome ^27^. To determine the role of Ca^2+^ in NLRP3 inflammasome activation in OCs, we assessed ASC speck formation and Sytox green uptake in *Tmem178^+/+^, Tmem178^−/−^*, and *Tmem178^−/−^*;*Nlrp3^−/−^* OC cultures exposed to exogenous CaCl_2_. As expected, LPS did not affect ASC speck formation and Sytox green uptake by OCs of any genotypes (Fig. 4B, D; Fig. S8A). CaCl_2_ stimulated ASC speck formation and Sytox green internalization in *Tmem178^−/−^*;*Nlrp3*^+/+^ OCs; it only promoted Sytox green uptake, but not ASC speck formation in *Tmem178*^+/+^;*Nlrp3*^+/+^ cells (Fig. 4B, D; Fig. S8). LPS + CaCl_2_ induced ASC speck formation (Fig. 4A). Sytox green internalization in response to these stimuli was higher in *Tmem178^−/−^* cultures compared to *Tmem178*^+/+^;*Nlrp3*^+/+^ and abrogated in *Tmem178^−/−^*;*Nlrp3^−/−^* OCs (Fig. 4A-D; Fig. S7B). These results suggest that Ca^2+^ acts as an important DAMP for the NLRP3 inflammasome in OCs. Consistent with this view and the notion that Tmem178 controls cytoplasmic Ca^2+^ influx, levels of intracellular calcium in this compartment were higher in *Tmem178^−/−^* cells compared to WT counterparts in response to LPS + CaCl_2_ or LPS + nigericin treatment (Fig. 4E). Furthermore, time-lapse imaging of multinucleated *Tmem178*^+/+^;*Nlrp3*^+/+^ OCs revealed higher mean fluorescence intensity of calbryte 520AM in OCs exposed to LPS + CaCl_2_ compared to LPS + nigericin, a transient response that was further induced in the latter group upon additional CaCl_2_ supplementation (Fig. 4F; Fig. S9 with video). Thus, in line with Tmem178’s function in restricting intracellular Ca^2+^ accumulation, Tmem178^−/−^ OCs exhibited significantly higher mean fluorescence intensity, which remained largely unchanged with additional CaCl_2_ supplementation.

**Fig. 4.**
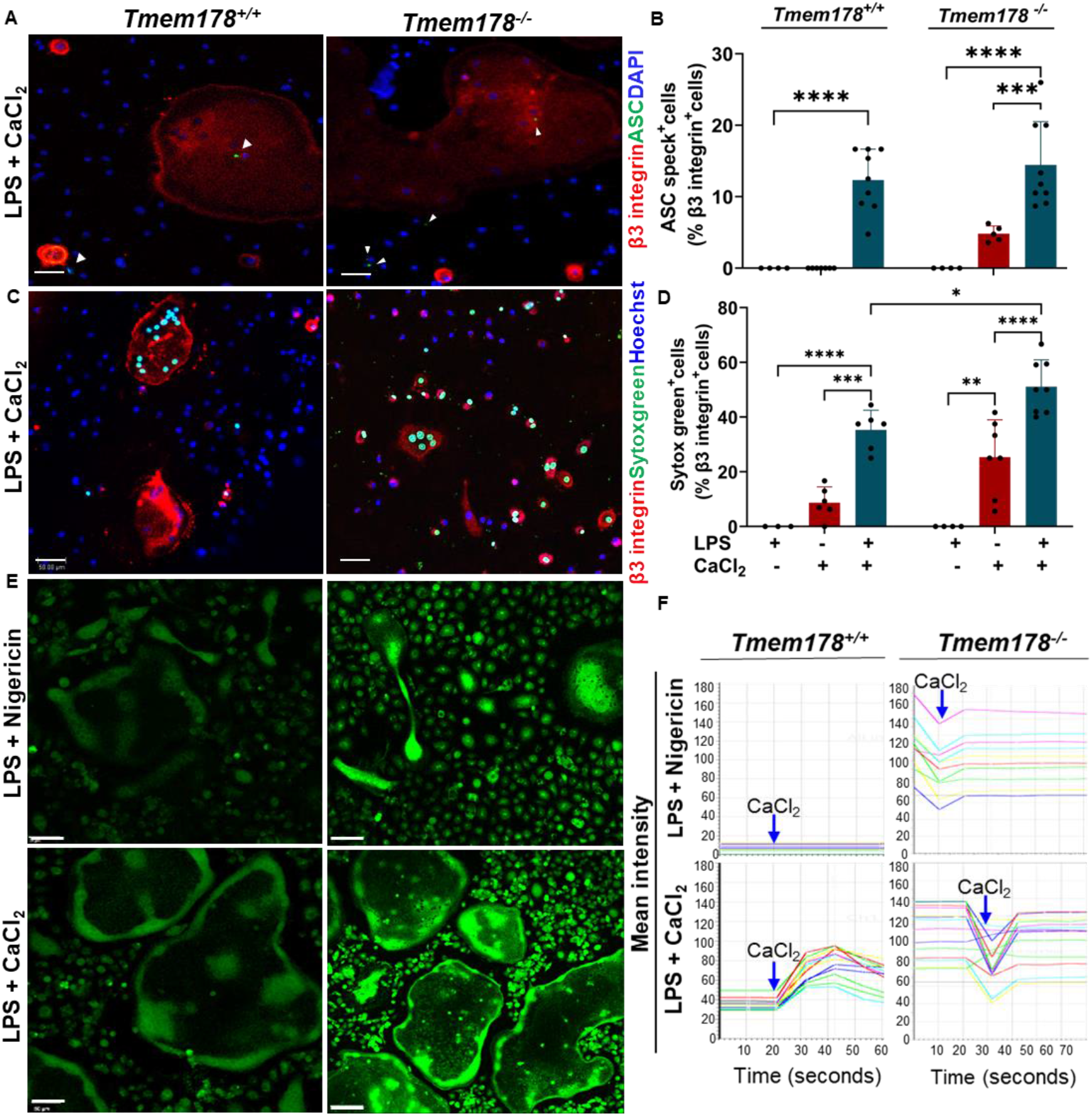
Loss of Tmem178 leads to high intracellular calcium levels and is associated with NLRP3 inflammasome activation. *Tmem178*^+/+^ and *Tmem178^−/−^* BMDMs treated with 100 ng/ml RANKL for 4-5 days were exposed to LPS for 3 h followed by 5 mM CaCl_2_ supplementation for 45 mins for ASC speck formation assay (A) and quantitative analysis (B). LPS-exposed cells were incubated with Hoechst and then with 5 mM CaCl_2_ for 45 mins for Sytox green uptake analysis (C) followed by β3 integrin immunostaining. Quantitative analysis of Sytox green uptake in β3 integrin positive OCs (D). The response to CaCl_2_ was measured using calbryte 520 AM, used at 5 µg/ml (final concentration) for 30 mins (E). Cells were stimulated again after 45 mins of treatment with 5 mM CaCl_2,_ and calcium flux was monitored using live time-lapse imaging (F). Each line represents the mean fluorescence intensity of calbryte 520AM of a multinucleated OC, which was in the 20-40 and 60-90 range, respectively, for *Tmem178*^+/+^ and *Tmem178^−/−^* OCs treated with LPS alone (F). Scale bar 50 µm. 2-way ANOVA with Tukey’s test, **p<0.005; ****p<0.0001.

### Ca^2+^ released from calcium-phosphate-coated plates activates the NLRP3 inflammasome

To investigate the effects of calcium release from the calcified bone matrix by OCs, we carried out cell cultures on plastic plates, calcium-phosphate-coated plates, which served as a surrogate for the bone matrix, and on bone slices. While Sytox green uptake and ASC speck formation induced by LPS and nigericin were confined to BMDMs on plastic plates (Fig. 1H; Fig. 5A), these responses occurred in both BMDMs and OCs on calcium-phosphate-coated plates (Fig. 5B, C).

**Fig. 5.**
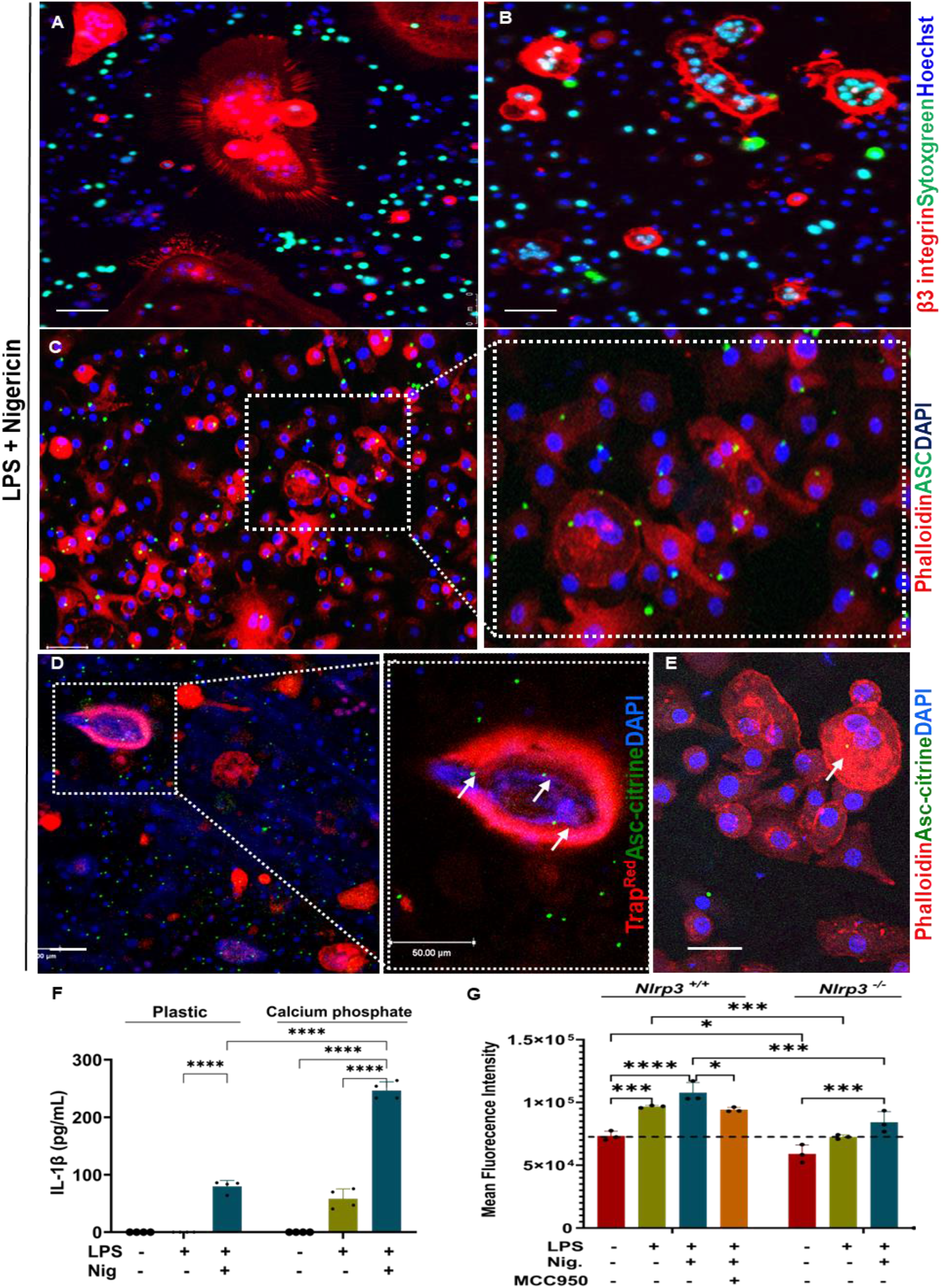
Bone-derived and CaP-released calcium triggers NLRP3 activation and enhances bone resorption. WT pre-OCs seeded on tissue culture (A) and calcium phosphate (CaP)-coated plates (B, C) were sequentially treated with RANKL for 3 days, LPS for 3 h, and nigericin for 45 min. Sytox green uptake was assessed using confocal microscopy on live cells (A, B). ASC speck formation was assessed in cells fixed after treatment and stained with anti-ASC antibody and phalloidin (C). To analyze OC activity, pre-OCs from *Trap*^*Red*^;*Asc-citrin*e mice were seeded on bone slices, treated with LPS and nigericin, and nuclei were stained with DAPI. Bone slices were visualized for ASC specks using a confocal microscope (D) and stained with phalloidin (E). IL-1β was measured in the culture supernatants (F). CaP plates were conjugated to fluorescein as per the kit protocol. *Nlrp3*^+/+^ and *Nlrp3^−/−^* OCs on conjugated plates were treated with LPS and/or MCC950 for 6 h, and nigericin for 1 h. Supernatants were collected to measure the release of fluorescein-conjugated matrix. Mean fluorescence intensity following an excitation wavelength of 485 nm and emission wavelength of 535 nm was measured using a spectrophotometer (G). Data analysis was done using 2-way ANOVA, Tukey’s test, *p<0.05; **p<0.01; ***p<0.005; ****p<0.0001.

Accordingly, TRAP-positive OCs differentiated from *Trap*^*Red*^; *Asc-citrine* pre-OCs seeded on bone slices, were deeply embedded in the bone surface and exclusively displayed ASC speck positivity (Fig. 5D; Fig. S10A-D with video). Multinucleated OCs displaying F-actin rings based on phalloidin staining were also ASC specks^+^ (Fig. 5E). Additionally, cells cultured on calcium-phosphate-coated plates exhibited significantly higher IL-1β secretion compared to those on plastic plates (Fig. 5F). Consistent with the idea that calcium could be released from calcium-phosphate-coated plates, LPS alone caused a notable increase in IL-1β levels in the supernatant (Fig. 5F).

To assess the impact of NLRP3 inflammasome activation on OC bone-resorbing activity, *Nlrp3*^+/+^ and *Nlrp3^−/−^* pre-OCs were incubated with RANKL for 3 days on fluorescein-conjugated calcium phosphate-coated plates. In addition, *Nlrp3*^+/+^ pre-OCs cells were pre-treated with MCC950, an NLRP3 inhibitor. The cells were then stimulated with LPS, either in absence or the presence of nigericin. LPS induced the release of fluorescein conjugated chondroitin sulfate bound to the calcium phosphate plate; this response was enhanced by nigericin and reduced by treatment with MCC950 or the loss of NLRP3 (Fig. 5G). These findings suggest that calcium influx is necessary and sufficient for NLRP3 inflammasome activation in OCs, and it enhances their bone resorbing activity.

### Bone loss caused by Tmem178 deficiency is prevented by the ablation of *Nlrp3*

Mice lacking Tmem178 exhibit a low bone mass phenotype associated with increased OC differentiation ^26^. The inhibition of NLRP3 inflammasome by Tmem178 provided a rationale for determining the impact of NLRP3 loss in osteopenia caused by Tmem178 deficiency. Micro-CT analysis revealed that bone microarchitecture was deteriorated in *Tmem178^−/−^*;*Nlrp3*^+/+^ mice, but not in *Tmem178*^+/+^; *Nlrp3*^−/−^ mice and *Tmem178*^+/+^;*Nlrp3*^+/+^ animals, a phenotype that was prevented in compound mutant counterparts (Fig. 6A). Accordingly, bone parameters, including bone mass (BV/TV), number of trabeculae (Tb.N), and bone mineral density (BMD) were slightly higher in *Tmem178*^+/+^;*Nlrp3*^−/−^ mice and *Tmem178^−/−^*;*Nlrp3^−/−^* mice compared to *Tmem178*^+/+^;*Nlrp3*^+/+^ controls, but were significantly reduced in *Tmem178^−/−^*;*Nlrp3*^+/+^ mice (Fig. 6B-D). Likewise, serum levels of cross-linked C-telopeptide type I collagen (CTX-I), biomarkers of bone resorption, were lower in *Tmem178^−/−^*;*Nlrp3^−/−^* mice compared to the other genotypes, among which CTX-I levels were similar (Fig. 6E). TRAP staining also revealed similar results, showing an increase in osteoclast number in *Tmem178^−/−^*;*Nlrp3*^+/+^ mice compared to another genotypes (Fig. 6F, G). These results suggest that the bone loss caused by the Tmem178 deficiency is, at least in part, mediated by the NLRP3 inflammasome.

**Fig. 6.**
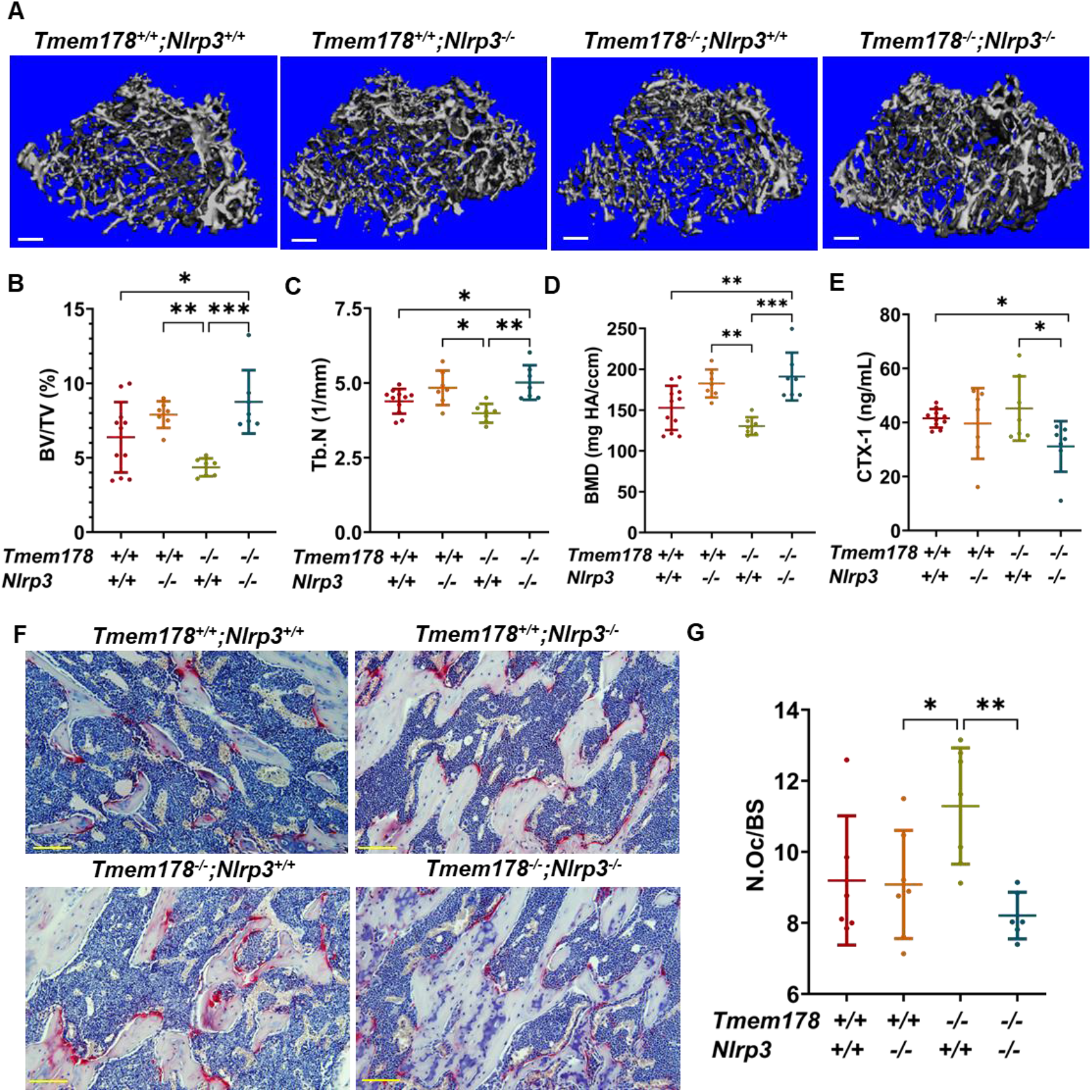
Bone loss caused by Tmem178 deficiency is prevented by the ablation of *Nlrp3*. Femurs from 10-week-old male *Tmem178*^+/+^;*Nlrp3^+/+^, Tmem178*^+/+^;*Nlrp3^−/−^, Tmem178^−/−^*;*Nlrp3*^+/+^, and *Tmem178^−/−^;Nlrp3^−/−^* mice were analyzed by micro-CT. Representative cross-sections from 3D reconstructions (A). Scale bar: 100µm. Quantification of bone parameters, including BV/TV, Tb.N, and BMD (B-D). Serum CTX-I levels were measured by ELISA (E). TRAP staining of histological bone sections showing representative images for each group (F). Scale bar: 50µm. Quantification of TRAP-positive OCs in the metaphyseal region, distal from the growth plate, shown as N.Oc/BS (G). Data are presented as means ± SD. Statistical significance was assessed using 1-way ANOVA. *p<0.05; **p<0.01; ***p<0.001. BMD, bone mineral density; BV/TV, bone volume/total volume; Tb.N, trabecular number; N.Oc/BS, OC number per bone surface.

## DISCUSSION

Under standard *in vitro* culture conditions, where cells were seeded on plastic, we found that the NLRP3 inflammasome could be activated in BMDMs but not in OCs. In contrast, the activation of this inflammasome was readily observed in these cells *in vivo*. To delineate the molecular mechanisms contributing to this discrepancy between *in vitro* and *in vivo* observations, we focused on Tmem178, a protein highly expressed in OCs and known to regulate Ca^2+^ flux, a damage-associated molecular pattern (DAMP) for the NLRP3 inflammasome ^28–31^. While the NLRP3 inflammasome was modestly activated in Tmem178-deficient macrophages, as previously reported ^35^, it was strongly activated in OCs lacking this protein. Several factors may account for these divergent outcomes: i) differences in Ca^2+^ signaling as these ions can act intracellularly and extracellularly through G protein-coupled receptors to activate the NLRP3 inflammasome ^29,36,37^; ii) the specific cellular context as various channels, including store-operated calcium channels and transient receptor potential channels, which mediate calcium internalization are not uniformly expressed across cell types ^35,38^; and iii) the dynamics of microstructures like actin filaments, which are regulated by Ca^2+^ and known to inhibit the NLRP3 inflammasome and are uniquely reorganized in functional OCs ^39,40^.

The prevailing dogma posits that the NLRP3 inflammasome complex in each activated cell is assembled at the centrosome, which also functions as the microtubule-organizing center ^41,42^. While single ASC specks per cell are typically observed, a few studies have shown the presence of multiple specks in a cell ^43,44^. In our study, multiple ASC specks of various sizes were readily detected in OCs *in vitro*. This observation is consistent with the reports indicating that multinucleated avian, human, and murine OCs maintained the individual centrosomes of the fused individual precursors ^45–47^. It remains unclear whether the activation of the NLRP3 inflammasome in OCs aims to perpetuate its bone-resorbing activity or to shorten its lifespan through pyroptosis. Our observation that ASC specks were prominent in OCs deeply embedded on bone surfaces (Fig. 5D; Fig. S10 with videos) suggests that the NLRP3 inflammasome may be activated to promote OC activity. This view is supported by the results indicating that loss of *Nlrp3* prevented the low bone mass phenotype in *Tmem178*-deficient mice.

Overall, changes in bone and OC parameters were more evident with the loss of Tmem178 than with NLRP3 deficiency. These findings align with prior studies showing similar baseline bone mass in WT and NLRP3 deficient young mice as well as the osteopenic phenotype observed in Tmem178 knockout mice compared to their littermate controls ^15,26^. Despite this difference, the impact of Tmem178 deficiency on the NLRP3 inflammasome and bone outcomes was normalized both *in vivo* and *in vitro* in the absence of NLRP3, suggesting that Tmem178 plays a crucial role in regulating the NLRP3 inflammasome.

In summary, we identified the Tmem178-calcium axis as a critical regulator of the NLRP3 inflammasome and bone resorption. Specifically, we found that Tmem178 serves as a negative regulator of the NLRP3 inflammasome in OCs, restraining bone resorption mediated by this pathway. Notably, this inhibitory effect can be overridden by environmental signals that induce the influx of calcium, which is released from bone by OCs. These findings suggest that the calcium-NLRP3 inflammasome pathway promotes OC activity and bone resorption as a positive feedback mechanism, which is negatively regulated by Tmem178.

## ACKNOWLEDGMENTS

Washington University Musculoskeletal Research Center (NIH P30 AR074992) supported Histology, Confocal microscopy, and analysis. Micro-CT imaging and analysis were supported by the Washington University Musculoskeletal Structure and Strength Core (NIH S10 OD028573).

We thank Crystal Idleburg, Samantha Coleman, and Michael Brodt for technical support.

## Funding

National Institute of Arthritis and Musculoskeletal and Skin Diseases, R01-AR076758, GM

National Institute of Allergy and Infectious Diseases, R01-AI161022, GM

National Institute on Aging, R01 AG077732, GM

National Institute of Arthritis and Musculoskeletal and Skin Diseases, R01-AR072623, YAA

National Institute of Arthritis and Musculoskeletal and Skin Diseases, R01-AR082192, YAA

Shriners Hospitals for Children, 85109, YAA

National Institute of Arthritis and Musculoskeletal and Skin Diseases, P30 AR074992, YAA

National Institute of Diabetes and Digestive and Kidney Diseases, T32 DK007120, NPS

The funders had no role in the study design, data collection, interpretation, or the decision to submit the work for publication

## Author contributions

Conceptualization: KK, YA, CW, GM

Methodology: KK, YA, CW, SB, KK, YL, NPS

Investigation: KK, CW, TP, NPS

Visualization: KK, CW

Funding acquisition: GM, YAA, DJV, RF

Project administration: GM

Supervision: CW, GM

Writing – original draft: KK, GM

Writing – review & editing: KK, GM, DJV, RF, YAA, NPS

## DATA AND MATERIALS AVAILABILITY

The study did not generate new, unique materials or reagents. All data are available in the main text or the supplementary materials.

## Lead contact

Further information and requests for resources and reagents should be directed to and will be fulfilled by the lead contact, Gabriel Mbalaviele.

## SUPPLEMENTARY MATERIALS

Supplementary Figure. Fig. S1-S10 Supplementary Table. Table S1

## MATERIALS AND METHODS

### Animals

Wild-type (WT) mice, *Asc-citrine*, and *Nlrp3^−/−^* mice were obtained from Jackson Laboratories. *Tmem178^−/−^* mice and *Trap* promoter-tdTomato (*Trap*^*Red*^) transgenic mice have been previously described (*26,34*). To generate *Tmem178^−/−^*; *Nlrp3^−/−^* and *Trap*^*Red*^; *Asc-citrine mice, Tmem178*^−/−^ were crossed with *Nlrp3^−/−^*, and *Trap*^*Red*^ mice were crossed with *Asc-citrine*, respectively. The resulting offspring were genotyped using primers specific for *Nlrp3, Tmem178, Trap^Red^*, and *Asc-citrine*. All mice were bred and housed in a pathogen-free environment at the Washington University School of Medicine animal facility. All mice were maintained under standard laboratory conditions, and experiments were conducted following institutional guidelines.

## Cell cultures

The femurs and tibiae were harvested from WT (*Tmem178^+/+^;Nlrp3*^+/+^), *Asc-citrine, Trap^Red^;Asc-citrine, Tmem178^−/−^;Nlrp3^+/+^, Tmem178^+/+^;Nlrp3^−/−^*, and *Tmem178^−/−^;Nlrp3^−/−^* mice were used to expand BMDMs *in vitro*. Bone marrow cells were flushed from bones as previously described ^48^. Cells were cultured in 10 mm Petri dishes in the presence of CMG (as a source of macrophage colony-stimulating factor, M-CSF) for 4-5 days. Expanded BMDMs were trypsinized and seeded into appropriate well plates at densities of 5 × 10^3^ cells per well in 96-well plates, 3 × 10^4^ cells per well in 24-well plates, and 0.4 × 10^6^ cells per well in 6-well plates. OC differentiation was induced by the addition of a receptor activator of nuclear factor kappa-B ligand (RANKL, 50-100 ng/mL) for 5-6 days. For calcium phosphate (CaP)-coated wells and bone slices, pre-OCs were seeded onto these substrata.

### RNA isolation and qRT-PCR

RNA was extracted from BMDMs and OC cultures using the PureLink RNA Mini Kit (Life Technologies) following the manufacturer’s instructions. DNA contamination was removed with PureLink DNase I (Invitrogen). The quality and quantity of RNA were assessed using a Nanodrop spectrophotometer (Thermo Scientific). cDNA was synthesized from 1 µg of RNA using the High-Capacity cDNA Reverse Transcription Kit (Applied Biosystems). Gene expression was analyzed by SYBR Green-based qRT-PCR using specific primers (Table S1). Data were normalized to cyclophilin B, and relative mRNA expression was calculated using the 2^^-ΔΔCt^ method.

### RNA-Seq

BMDMs were cultured for 5 days in 2% CMG, then left untreated or treated with RANKL to generate OCs. On day 5, cells were treated with or without 100 ng/mL LPS for 3 h. RNA was extracted using the PureLink RNA Mini Kit (Invitrogen) with on-column DNase treatment (Ambion). RNA quality and quantity were assessed using the Agilent Bioanalyzer 2100. Libraries were prepared using the NEBNext® Ultra™ RNA Library Prep Kit and sequenced on an Illumina platform to generate paired end reads. Raw reads were cleaned using fastp and aligned to the reference genome, mm10 using STAR. Gene expression was quantified with Feature Counts, and RPKM values were calculated. Differential expression was analyzed using DESeq2 with Benjamini-Hochberg correction (adjusted p<0.05). Three biological replicates were used per group. Raw and processed RNA-seq data are publicly available in the NCBI GEO database under accession number, GSE298028.

### Western blot analysis

Cellular extracts were prepared by lysing cells with RIPA buffer (50 mM Tris, 150 mM NaCl, 1 mM EDTA, 0.5% NaDOAc, 0.1% SDS, 1.0% NP-40) containing protease inhibitors (GenDEPOT, TX). Protein concentrations were measured using the Bio-Rad method (Bio-Rad, CA), and 30-40µg of protein were separated on 12% SDS-PAGE gels as described by Wang et al., 2024. Proteins were transferred to nitrocellulose membranes and incubated overnight at 4°C with antibodies against GSDMD (1:1000; Abcam, MA), NLRP3 (1:1000; AdipoGen, CA), β-integrin (1:1000; Cell Signaling Technologies, MA), and β-actin (1:2000; Santa Cruz Biotechnology, TX). Afterward, membranes were incubated with secondary antibodies and visualized using the Odyssey infrared imaging system (LI-COR Biosciences, NE).

### Sytox green uptake assay

LPS- or TNF-α-primed BMDM and OC cultures were incubated with Hoechst 3642 (1 µg/mL) for 15-20 mins, fed with fresh media, then treated with 15 µM of nigericin or 5 mM calcium chloride (CaCl_2_) for 45 mins. Cells were incubated for 10-15 mins with Sytox green (Invitrogen) diluted in PBS at a final concentration of 10 nM and stained with an Alexa Fluor 647-labeled β3 integrin antibody (1:500; BD Pharmingen). Fluorescence in live cells was measured using a Leica DMi8 confocal microscope. Sytox green- or β3 integrin-positive cells were counted using ImageJ.

### ASC speck formation assay

To analyze ASC speck formation *in vitro*, LPS– or TNFα–primed cells were stimulated with nigericin or CaCl_2_, followed by blocking and staining for β3-integrin (Alexa Fluor 647-labeled β3-integrin antibody) for 30 min at 4°C. After staining, the cells were washed and fixed with 4% PFA for 10 mins, then permeabilized and stained for ASC using an anti-mouse ASC antibody (EMD Millipore) diluted in antibody buffer (0.2% Triton X-100 and 1% BSA in PBS) and incubated overnight at 4°C. The following day, cells were washed and incubated with an Alexa Fluor 488-conjugated anti-mouse IgG secondary antibody (Invitrogen) and mounted with DAPI-containing mounting medium. ASC speck formation was detected using immunofluorescence microscopy. The number of ASC speck^+^ and/or β3-integrin^+^ cells was analyzed using ImageJ.

To assess ASC speck formation *in vivo, Trap^Red^; Asc-citrine* reporter mice were injected subcutaneously with RANKL (2 mg/kg body weight) over the periosteum of the calvaria for four consecutive days. On day 5, mice were administered 15 mg/kg LPS for 6 h. Calvariae were harvested, fixed in 4% PFA at room temperature for 8 h and then overnight at 4°C. The tissues were washed multiple times in PBS, decalcified in 14% EDTA for 3 days, infiltrated with 30% sucrose overnight at 4°C, embedded in OCT, and sectioned into 5-μm slices. Tissue sections were stained with DAPI, and ASC speck formation was examined using a Leica DMi8 microscope, with lasers set at 405 nm, 488 nm, and 561 nm to excite DAPI, citrine, and tdTomato fluorescence, respectively.

### Cytokine release assay

Culture supernatants were collected after LPS priming and nigericin stimulation, and IL-1β levels were quantified using an ELISA kit (eBiosciences) following the manufacturer’s instructions. Cell death was measured by the release of lactate dehydrogenase (LDH) using an LDH cytotoxicity assay kit (Roche).

### Intracellular calcium flux assay

To analyze intracellular calcium fluxes, OC cultures were primed with LPS for 3 h, followed by incubation with 1X Calbryte 520 AM solution at a final concentration of 2.5 µM for 30 min. The cells were then washed with 1X HBSS (Hank’s Balanced Salt Solution), and fresh media were added. Subsequently, the cells were stimulated with either nigericin or CaCl_2_ for 45 mins, after which fluorescence was measured using confocal microscopy. Time-lapse imaging was performed during the addition of (nigericin or CaCl_2_) to the LPS-primed cells using a Leica DMi8 microscope.

### Bone resorption assay

Bone resorption was measured using fluorescein amine-coated CaP plates (Cosmo Bio, USA). The binding of fluorescein amine-labeled chondroitin sulfate (FACS) to the CaP-coated plates was carried out before cell seeding, following the manufacturer’s instructions under sterile conditions. This was done by adding 100 µL of bone resorption assay FACS solution to the plate, which was then incubated for 3 h at 37°C, followed by 2 washes with PBS. Plates were kept in light-shielded conditions. Pre-OCs treated with RANKL for 2-3 days were seeded onto the coated CaP plate in the phenol-red-free medium. The cells were incubated for an additional 3 days in the presence of RANKL without changing the medium. On day 4, the cells were treated with LPS and/or MCC950, a selective NLRP3 inhibitor, for 6 h, followed by stimulation with nigericin for 1 h. After stimulation, 100 µL of medium from each well was transferred to a 96-well black plate for fluorescence measurements, and 50 µL of assay buffer was added to each well before measuring fluorescence. Fluorescence intensity was measured by exciting the cells at a wavelength of 485 nm and recording the emission at 535 nm. Media without cells served as the blank.

### Bone mass analysis using µCT

Bone mass and microstructure of 10-week-old male mice were evaluated using a Scanco Micro-CT50 scanner. Femora were cleaned of soft tissues, stabilized in 2% agarose gel, and scanned at 10 μm resolution. Approximately 150 slices of the distal femoral metaphysis were analyzed for bone parameters. The trabecular bone region was manually traced, threshold at 600, and reconstructed into 3D images using Scanco analyzer software, as previously described ^48^.

### Measurement of Bone Resorption Using CTX-I Assay

To assess bone resorption, C-terminal telopeptides of type I collagen (CTX-I) were measured in mouse serum. Serum samples were collected from 10-week-old male mice after a minimum 6 h fasting period. The levels of CTX-I were quantified using the RatLaps® (CTX-I) EIA kit (RatLaps™, IDS, Boldon, UK) following the manufacturer’s protocol.

### Histology

To detect OCs, bones were fixed overnight in 10% buffered formalin and decalcified in 14% EDTA (pH 7.0) for 10 days (long bones) or 3 days (calvaria) with gentle rocking and daily solution changes. Decalcified bones were dehydrated in graded alcohol, cleared with xylene, and embedded in paraffin (FFPE). Longitudinal sections (5 μm) were stained with TRAP (Sigma) to identify OCs and counterstained with hematoxylin. Imaging was performed using a Nikon Eclipse 80i microscope with a 20X objective.

### Statistical analysis

All the **s**tatistical analysis was carried out using GraphPad Prism 9.0 software. Methods included the student’s t-test, one-way ANOVA with Tukey’s multiple comparisons test, and two-way ANOVA with Dunnett’s multiple comparisons test. Results are shown as mean ± SD, with *p* < 0.05 indicating statistical significance.

