## Supplemental figures S1-S10, Supplemental table S1 for "NLRP3 inflammasome is regulated in osteoclasts through a Tmem178-dependent mechanism that restricts calcium influx"

**Supplementary Data**

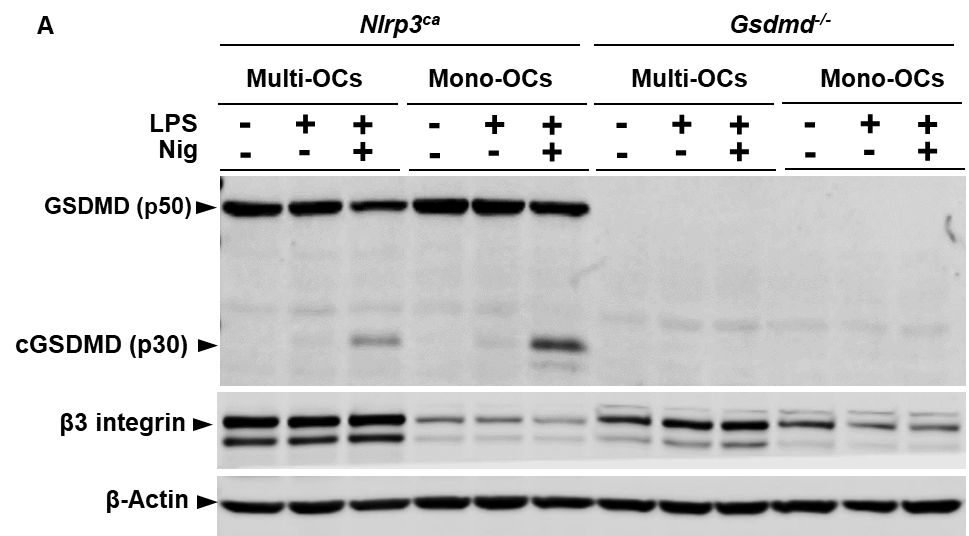

**Fig. S1. NLRP3 inflammasome-mediated cleavage of GSDMD is lower in multinucleated OCs than mononucleated cells.** BMDMs from *Nlrp3^ca^* and *Gsdmd^-/-^* mice were treated with RANKL for 4 days, then exposed to LPS for 3 h and nigericin for 45 mins. Cell lysates were immunoblotted for GSDMD, cleaved GSDMD (cGSDMD), β3 integrin, and β-Actin.

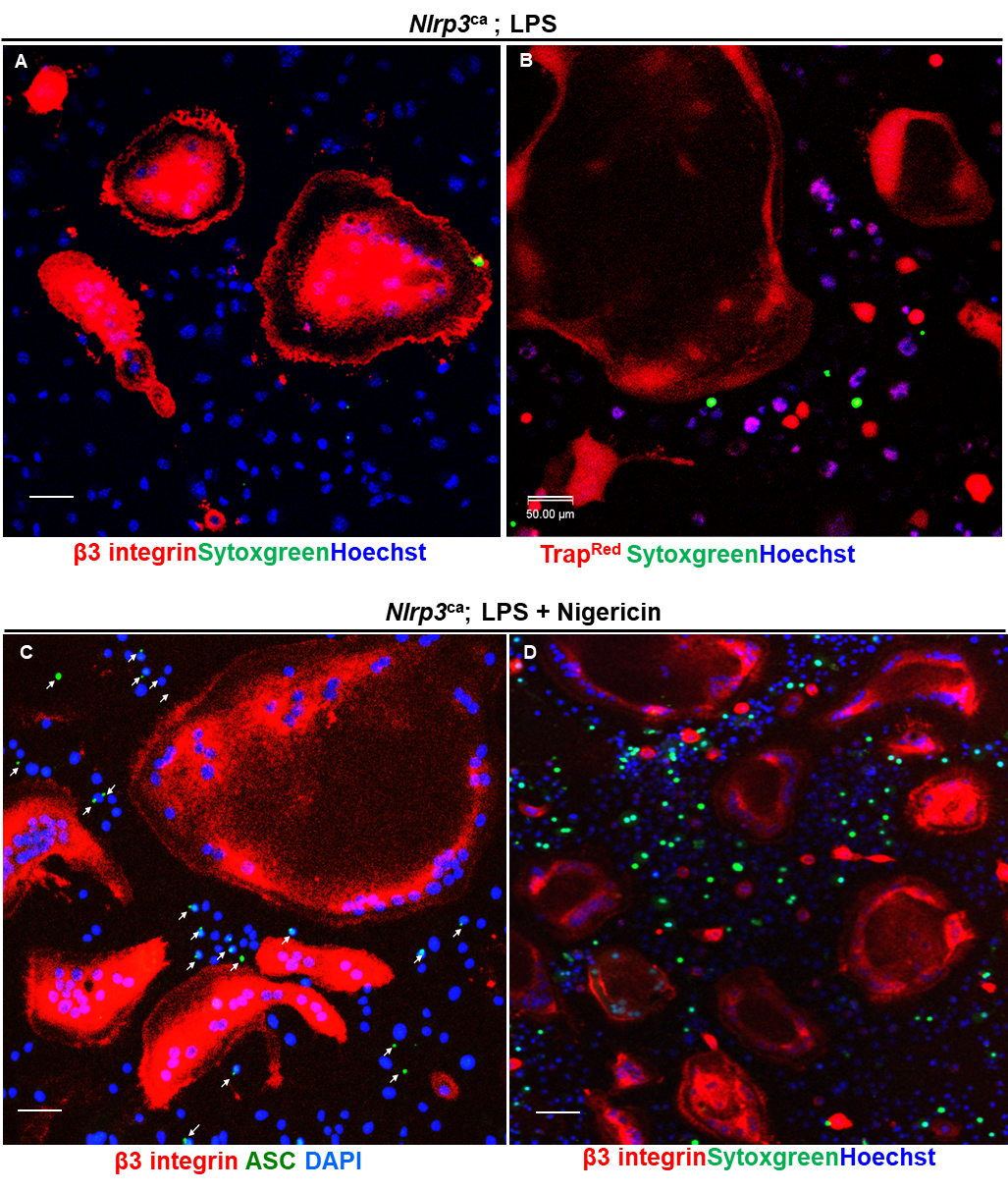

**Fig. S2. The inflammasome assembled by NLRP3^ca^ is impaired in the OC lineage:** NLRP3^ca^ OC cultures were exposed to LPS for 3 h (A, B), and then with nigericin for 45 mins (C, D) to analyze Sytox green uptake (A, B, D) and ASC speck formation (C). Cells were stained with anti-ASC antibody (green), β3 integrin antibody (red), and counterstained with DAPI (blue). White arrowheads indicate ASC specks (C). Scale bar 50 µm

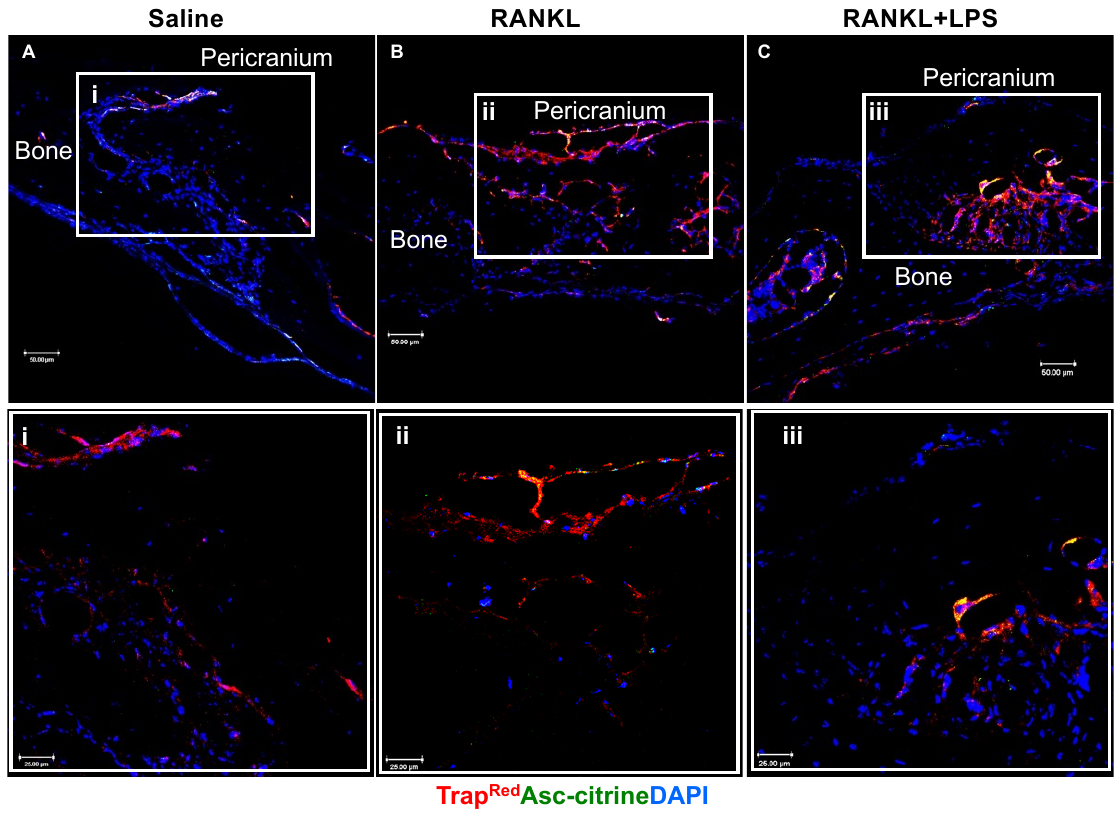

**Fig. S3. OCs assemble NLRP3 inflammasome *in vivo.*** Calvarial sections from *Trap^Red^*; *Asc-citrine* mice were treated with saline (A, i), RANKL (B, ii), or RANKL + LPS (C, iii). Images were taken at 20X magnification (A-C) and 40X, scale bar 50 µm and 25 µm respectively.

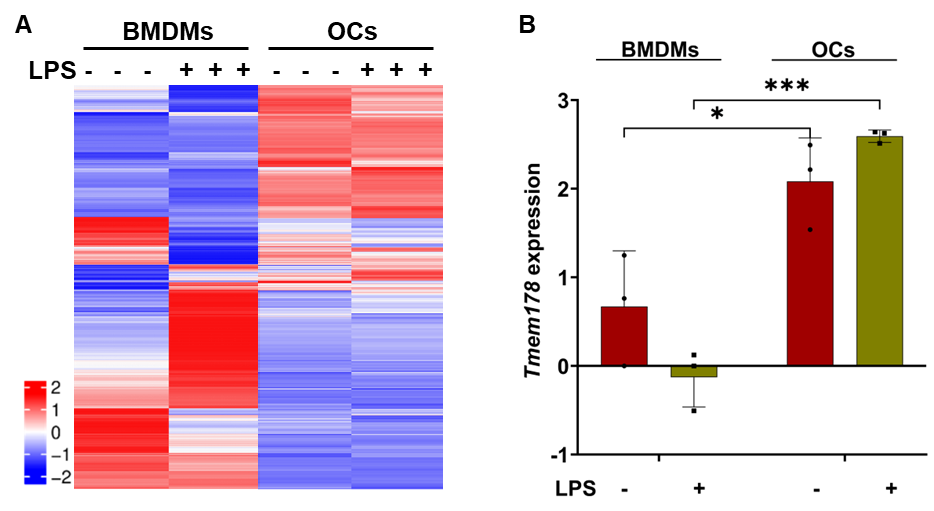

**Fig. S4. Transcriptomic profiling of BMDMs and OCs reveals differential gene expression and upregulation of *Tmem178*.** Heat map showing differentially expressed genes (DEGs) identified by RNA-seq in BMDMs and RANKL-treated BMDMs (OCs) left untreated or treated with 100 ng/mL LPS for 3 h. Three biological replicates per condition were analyzed (A). Expression levels of *Tmem178* across conditions, showing significantly higher expression in OCs calculated using DESeq2 with Benjamini-Hochberg correction (B).

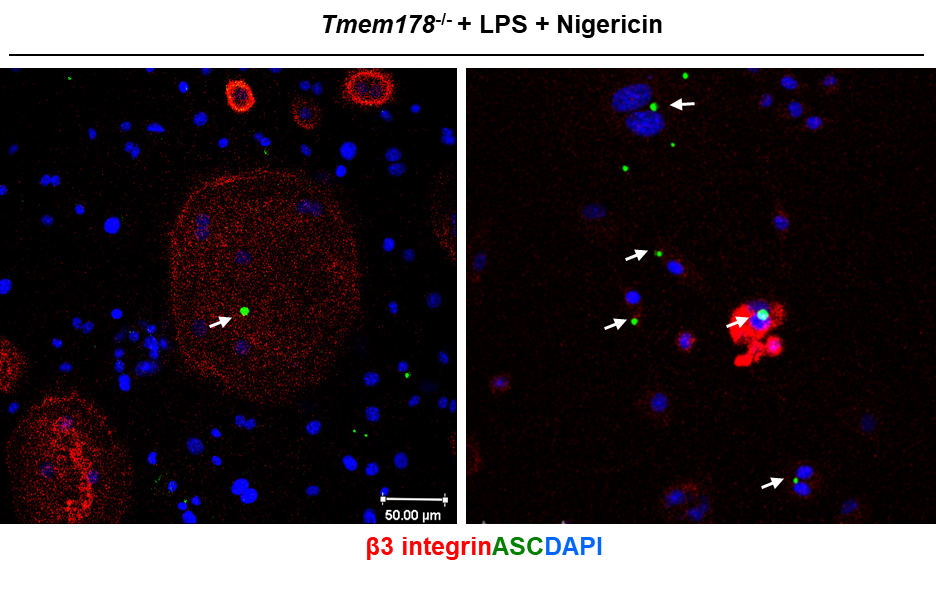
**Fig. S5. Tmem178 inhibits the activation of the NLRP3 inflammasome in the OC lineage.** *Tmem178^-/-^* BMDMs treated with RANKL for 5 days were exposed to LPS and nigericin. Cells were incubated with ASC antibody (green), β3 integrin antibody (red), and counterstained with DAPI (blue). White arrowheads indicate ASC specks in β3 integrin-stained OCs.

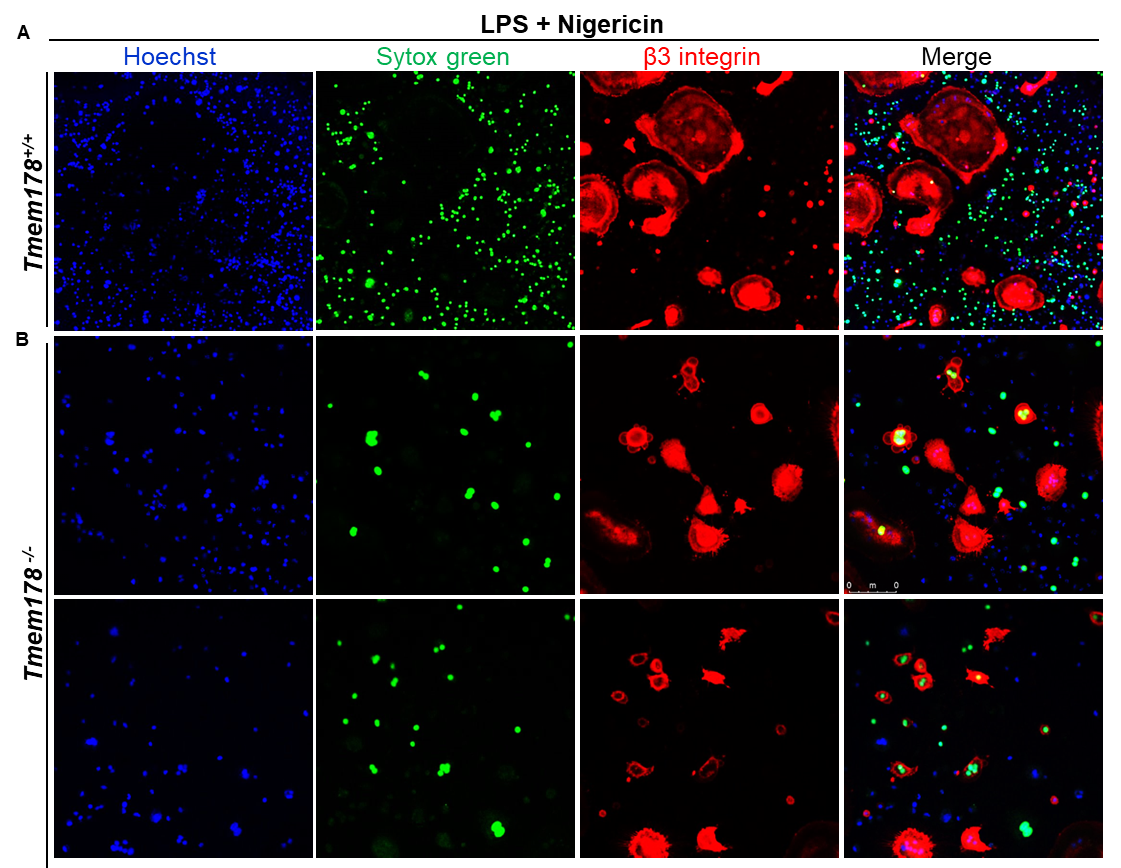

**Fig. S6: *Tmem178^-/-^* but not *Tmem178^+/+^* OCs internalize Sytox green.** *Tmem178^+/+^* (A) and *Tmem178^-/-^* (B) cells were exposed to LPS for 3 h then to Hoechst for 15 mins followed by nigericin (Nig) and Sytox green (green) for 45 mins. Cells were stained with β3 integrin antibody (red), and live cells were imaged using a confocal microscope—scale bar 50 µm.

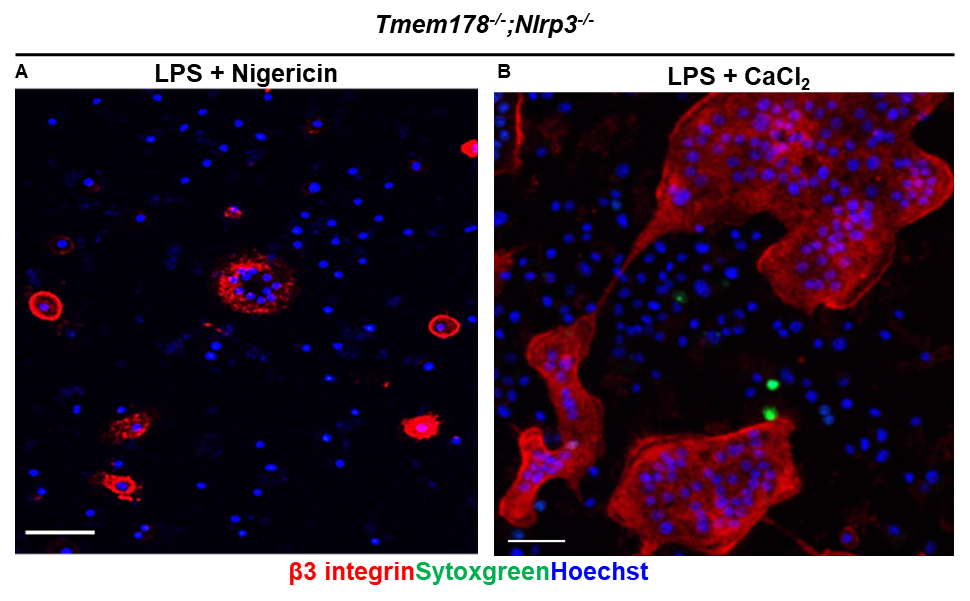
**Fig. S7: Tmem178 functions upstream of NLRP3:** *Tmem178^-/-^*;*Nlrp3^-/-^* OC cultures were exposed to LPS for 3 h, then with Hoechst for 15 mins followed by nigericin (A) or CaCl_2_ (B) for 45 mins, and incubated with Sytox green for 15 mins—scale bar 50 µm.

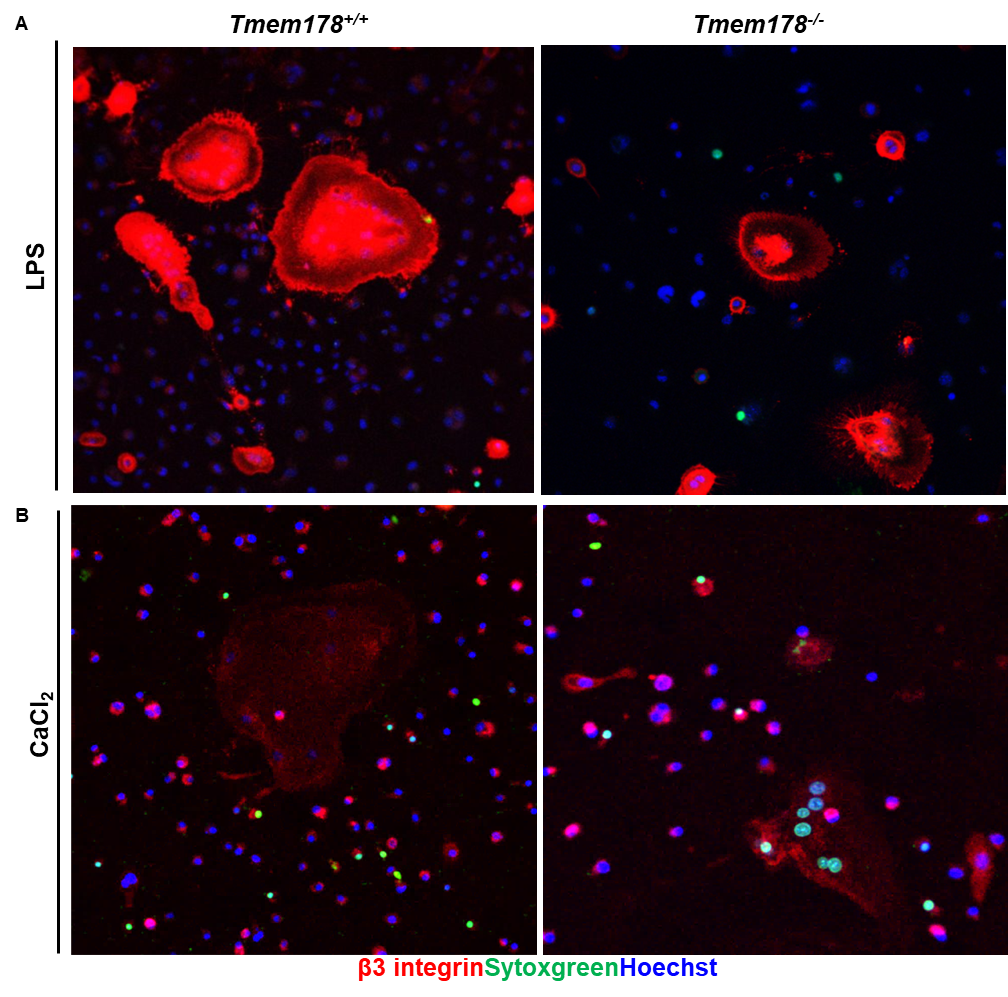

**Fig. S8: LPS or CaCl_2_ alone does not activate the NLRP3 inflammasome in OCs.** *Tmem178^+/+^* (A) and *Tmem178^-/-^* (B) OC cultures were exposed to LPS for 3 h (A) or CaCl_2_ for 45 mins (B), then with Hoechst (blue) for 15 mins and Sytox green (green) for 45 mins. Cells were stained with β3 integrin antibody (red) and live cells imaged using confocal microscope. Scale bar 50 µm.

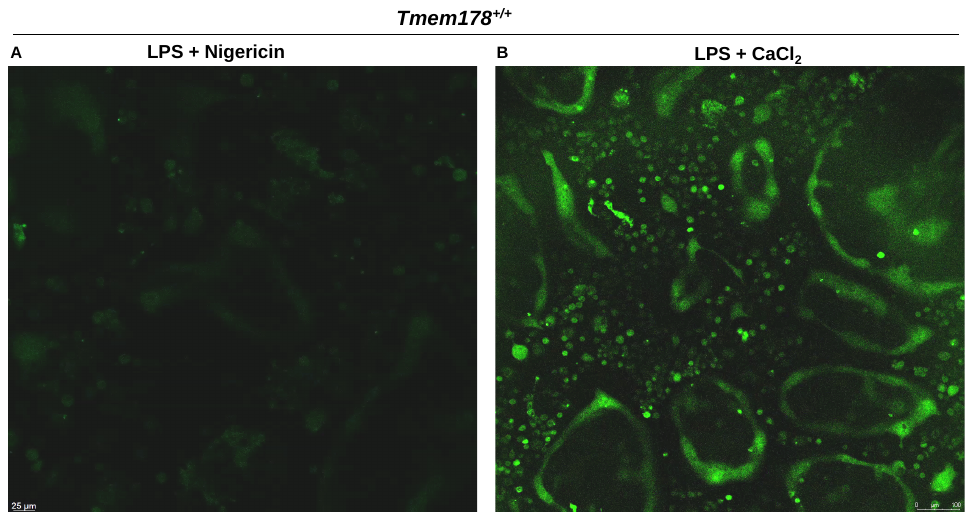
**Fig. S9 (video): Exogenous CaCl_2_ induces calcium influxes in OCs:** *Tmem178^-/-^* OC cultures were exposed to LPS for 3 h followed by incubation with 5 µg/ml calbryte 520 AM for 30 mins and nigericin (A) or CaCl_2_ for 45 mins (B)_._ Cells were stimulated again at the time of imaging with CaCl_2,_ and calcium flux was monitored using a time-lapse imaging.

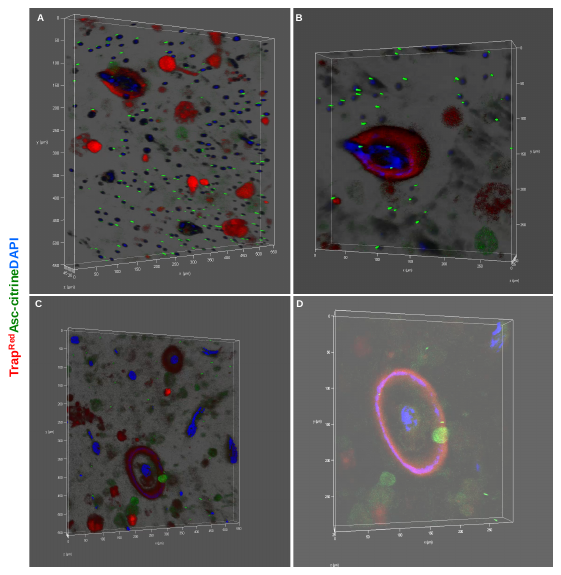

**Fig. S10-video. Deeply seated OCs on bone are ASC specks^+^.** Pre-OCs from *Trap^Red^*; *Asc-citrine* mice were seeded onto bone slices and treated with RANKL exposure for 5 days, followed by LPS for 3 h, and nigericin for 1 h. Cells were stained with DAPI. Bone slices were imaged using a confocal microscope with Z-sections up to a depth of 40 µm. 3D images were generated using LAX software.

**Supplementary Table**

**Table S1:** qPCR Primer List

| *Cyclophilin B* Fwd | AGCATACAGGTCCTGGCATC |
| --- | --- |
| *Cyclophilin B* Rev | TTCACCTTCCCAAAGACCAC |
| *Nlrp3* Fwd | TGCTCTTCACTGCTATCAAGCCCT |
| *Nlrp3* Rev | ACAAGCCTTTGCTCCAGACCCTAT |
| *Tmem178* Fwd | ATGACAGGGATATTTTGCACCAT |
| *Tmem178* Rev | CCGGTTCAAGTCATAGGAGACACT |
